# Hippo pathway perturbation disrupts cell fate control in the *Drosophila* eye

**DOI:** 10.64898/2026.05.08.723765

**Authors:** Abdul Jabbar Saiful Hilmi, Samuel. A Manning, Lucas. G Dent, Katrina A. Mitchell, Kieran F. Harvey

**Author notes:** London Centre for Nanotechnology, University College London, London, UK WC1E 6BT. Department of Biochemistry and Pharmacology, The University of Melbourne, Parkville, Victoria, Australia, 3010 (Katrina A. Mitchell).

## Abstract

During animal development, cells acquire specialised fates in a precise spatiotemporal order, which is essential for producing tissues that function appropriately. Cell fate specification is governed by multiple signalling pathways, as well as mechanical forces, which impact cellular transcription. Two such signalling pathways are the Hippo pathway and EGFR pathway, which both control organ growth and the fate of certain cell types in multiple species. Here, we show that Hippo signalling is essential for the maintenance of the cone and primary pigment cell fates in the developing *Drosophila* eye. When Hippo signalling is compromised, its nuclear effectors Yorkie and Scalloped drive increased expression of the EGFR pathway transcription repressor Yan, which antagonises the cone and primary pigment cell fates. Thus, in addition to its role as a growth suppressor, Hippo signalling promotes the fate of multiple eye cells by maintaining their responsiveness to inductive cues from the EGFR pathway.

**AUTHOR SUMMARY:** As multicellular organisms grow and develop from a zygote, individual cells become increasingly specialised. Cell fate is specified and maintained by the coordinate action of different signalling pathways, whose activity must be tightly controlled in a spatiotemporal fashion. If this fails, cell fate can be disturbed, which can cause developmental abnormalities, and diseases such as cancers. Here, we describe a new function for the Hippo signalling pathway, which was originally discovered as a regulator of *Drosophila* tissue growth and subsequently linked to the genesis of multiple human cancers. Hippo signalling is essential for maintaining the fate of two key cell types in the *Drosophila* eye, primary pigment cells and cone cells. Without Hippo signalling these cells cannot properly respond to signals from another key signalling network, the EGFR pathway. Our discoveries add to a growing literature where the Hippo growth control pathway is repurposed to control cell fate in tissues that have ceased growth.

## INTRODUCTION

The *Drosophila* compound eye is composed of about 800 eye units called ommatidia [1]. Each individual ommatidium consists of various cell types, with each of these undergoing a specific, tightly controlled, differentiation program [2, 3]. At the apical core of each ommatidium are four cone cells, which are surrounded by two primary pigment cells, and packed in a hexagonal shape by the interlacing of the secondary interommatidial cells, tertiary interommatidial cells, and the bristle complex cells [4, 5]. The deeper layers of each ommatidium also contain eight photoreceptor neurons, which sense visual cues and transmit them to the brain [1, 5, 6] (**Fig. 1A-D**). The different cell types of the *Drosophila* eye are readily detectable at the early pupal stage, with most cells adopting characteristic shapes and positions in the epithelium by 24 hours after puparium formation (APF) [2]. Cell nuclei also adopt specific positions in the eye; at the apical most layer are the cone cell nuclei, followed by the photoreceptor neurons, which reside alongside the nuclei of the two primary pigment cells. The nuclei of the interommatidial cells and bristle complex cells reside towards the basal side of the eye (**Fig. 1A-D**).

**Figure 1.**
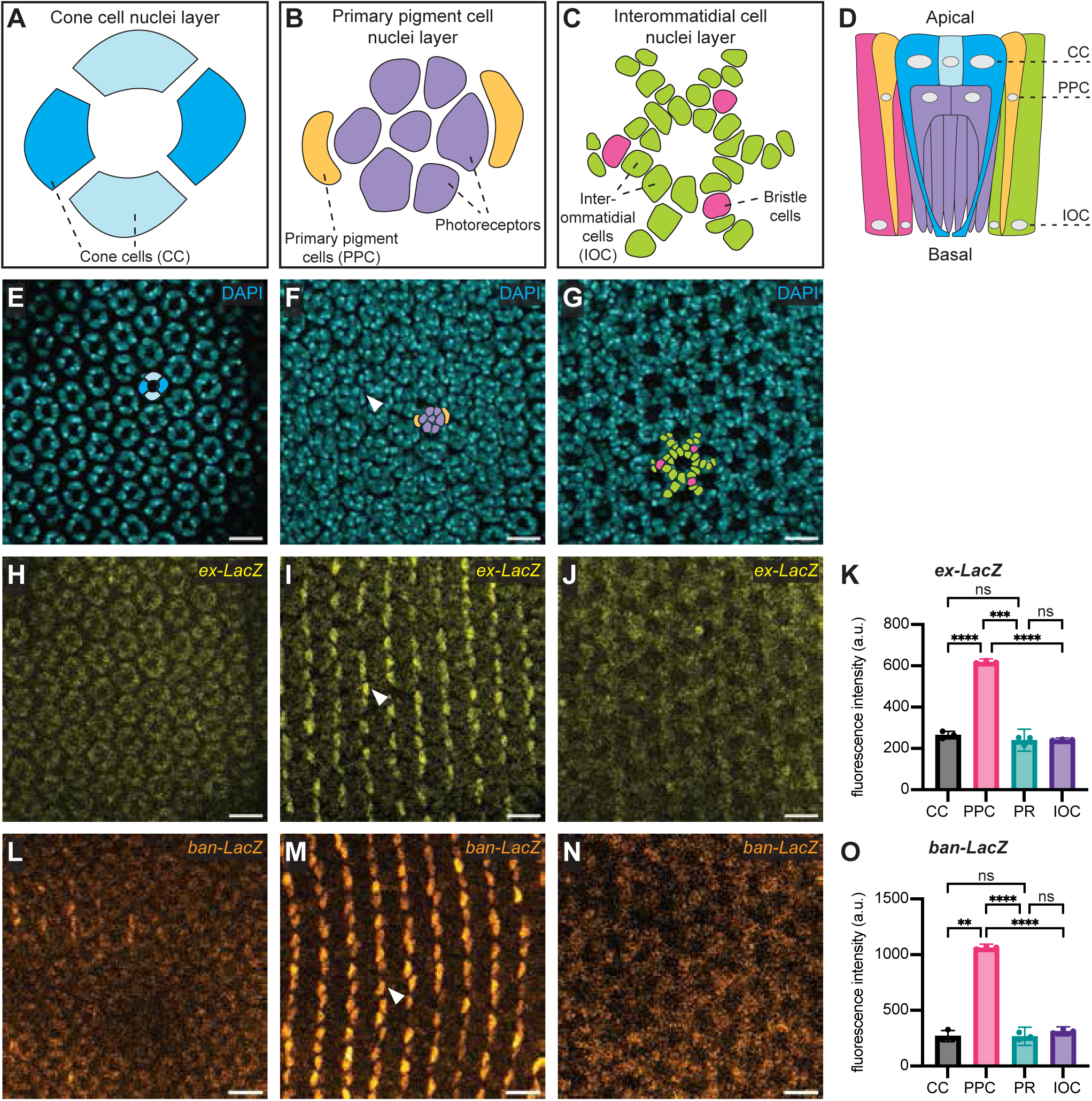
Yorkie transcription activity is elevated in primary pigment cells of the pupal *Drosophila* eye. (A-D) Schematic diagrams of the *Drosophila* pupal eye cell types. Cross-sections of the pupal eye along the X-Y axis at the nuclei layer of cone cells (A), primary pigment cells (B) and interommatidial cells (C). A cross-section of the pupal eye along the X-Z axis is depicted in (D). Cell types are represented in blue (cone cells), yellow (primary pigment cells), photoreceptor cells (purple), interommatidial cells (green), and bristle group cells (pink). Nuclei are indicated in grey. Adapted from [2]. (**E-J and L-N**) Confocal microscope images of a *Drosophila* pupal eye 24h APF at the cone cell nuclei layer (E, H, L), the primary pigment cell nuclei layer (F, I, M), and the interommatidial cell nuclei layer (G, J, N). DAPI (cyan) marks nuclei (E-G), *ex-LacZ* is in yellow (H-J) and *ban-LacZ* is in orange (L-N). In (E-G), schematic diagrams highlight nuclei of different cell types, with different colours representing the cell types in (A-D). Arrowheads indicate select primary pigment cells. Scale bars = 10μm. (**K and O**) Charts showing quantification of *ex-LacZ* (K) and *ban-LacZ* (O) fluorescence intensity in cone cells (CC), primary pigment cells (PPC), photoreceptor cells (PR), and interommatidial cells (IOC). n=3 for each sample, *** indicates P-value<0.0005, **** indicates P-value<0.00005, and ns indicates P-value>0.05, assessed by multiple comparisons test after ANOVA, with corrections applied for multiple comparisons. Interval bars indicate standard deviation.

Eye cell fate specification begins in the third instar period of larval development when differentiating photoreceptor cells emerge in regularly spaced rows, beginning with the R8 cells, which recruit other photoreceptor cells in a sequential manner [6–8]. Subsequently, during the late stages of the third larval instar, the non-neuronal cells begin to be specified, commencing with the cone cells [2]. Finally, during the pupal stage of development, the pigment and bristle complex cells emerge [2, 5, 9]. Cone cell development requires the transcription factor DPax2, the transcription of which is regulated by combined inputs of Notch and EGFR signalling from the neighbouring photoreceptors [7, 10–12]. Notch signalling promotes the activity of its downstream transcription factor Suppressor of Hairless [Su(H)], while EGFR signalling promotes the activity of the Pointed (Pnt) transcription factor, both of which bind to the enhancer region of *dPax2* at multiple sites [11, 13]. EGFR signalling further promotes *dPax2* transcription by repressing the transcriptional repressor Yan (also known as Anterior open, Aop) by phosphorylation and nuclear exclusion [11]. The transcription of *dPax2* is also promoted by the transcription factor Lozenge (Lz) [14], which is itself activated by the Sine oculis and Glass transcription factors [15]. Interestingly, Lz also acts as a transcriptional repressor together with Groucho, to repress the neuronal gene *deadpan* (*dpn*), thereby reinforcing the non-neuronal cone cell fate [16]. However, the combined inputs from Lz, Pnt, and Su(H) alone are not sufficient to drive cone cell differentiation, suggesting that other factors combine to regulate cone cell fate [13].

The Hippo pathway is a central signalling pathway that was first discovered as a regulator of tissue growth in genetic screens conducted in the *Drosophila* eye [17–25]. Unlike most signalling pathways, which are regulated by ligand-receptor protein pairs, the Hippo pathway responds primarily to mechanical cues like stretch and compression [26–32]. Hippo signalling controls transcription by phosphorylating and repressing the activity of the Yorkie (Yki) transcription co-activator (YAP/TAZ in mammals) [33–36]. This limits Yki nuclear abundance as well as the chromatin association times of both Yki and its DNA binding partner Scalloped (Sd) (TEAD1-4 in mammals, respectively) [33–42]. Perturbations in Hippo signalling cause eye overgrowth because larval eye cells proliferate faster and then fail to exit the cell cycle in a timely fashion [17, 18, 21–25]. These phenotypes are compounded by the fact that apoptosis that normally removes excess cells in the mid-pupal stage of eye development is defective in tissue lacking core Hippo pathway proteins like Salvador (Sav), Warts (Wts) and Hippo (Hpo) [17, 18, 21, 22, 24, 25].

Subsequently, Hippo signalling was also found to control the fate of a specific subset of *Drosophila* eye cells, i.e. during late pupal eye development, it dictates whether R8 photoreceptor cells adopt the pale or yellow fate to detect different wavelengths of light [43, 44]. These functions of the Hippo pathway are conserved in mammals, as its disruption causes the overgrowth of many tissues, whilst it is required for select cell fate choices, most notably the first cell fate decision in mammals, i.e. trophectoderm versus inner cell mass [45, 46]. However, genetic analyses of larval *Drosophila* eyes lacking *sav*, *hpo* or *wts* indicated that Hippo signalling is dispensable for the induction of cell fate during larval development [17, 18, 21, 23, 24, 47]. Potential roles for Hippo signalling in eye cell fate maintenance and cell fates that are induced in pupal development like pigment and bristle cells have not been explored thoroughly. Here, we show that Hippo signalling is required to promote the maintenance of both cone cells and primary pigment cells in the *Drosophila* eye. Hippo signalling does this by influencing the transcriptional output of the EGFR signalling pathway. Specifically, when Hippo signalling is disabled, Yki is hyperactivated and induces expression of the Yan transcription repressor, which can antagonise cone and primary pigment cell fate by regulating transcription of the key cell fate inducer DPax2.

## RESULTS

### Yorkie transcription activity is elevated in primary pigment cells of the pupal *Drosophila* eye

During the larval phase of *Drosophila* development, the eye, which initially consists of a pool of undifferentiated progenitor cells, grows substantially before commencing differentiation during the third larval instar [2, 3]. The larval eye is comprised of two epithelial sheets: the disc proper, which is a columnar epithelium, and the peripodial epithelium, which is squamous. The disc proper is the major contributor to the presumptive adult eye, giving rise to the light-sensing photoreceptor neurons, as well as cone cells, pigment cells and bristle complex cells [2, 3]. In the third instar larval eye disc, Yki/Sd activity is relatively even within each epithelial layer, although it is higher in the peripodial epithelium than the disc proper [48–50]. This suggests that Hippo signalling is also relatively even during the larval growth phase of the eye as it shows no major patterns of activity (**Fig. S1A**), in contrast with other signalling pathways like Hedgehog and EGFR, which display defined activity patterns during *Drosophila* larval eye development [3, 51]. To investigate whether Hippo pathway activity is uniform or patterned in the postmitotic phase of eye development, we assessed Yki/Sd-dependent transcription in the pupal eye, 24h after puparium formation (APF). At this developmental stage, proliferation has ceased, all major cell types have been specified and supernumerary interommatidial cells are yet to be removed by apoptosis. Strikingly, activity of the well-defined Yki/Sd transcription reporter *expanded-LacZ (ex-lacZ)* was non-uniform; instead, it was considerably higher in some cells compared to others, in regularly spaced rows (**Fig. 1H-J**). By defining the apicobasal location and distinctive shapes of cell nuclei within the pupal eye by DAPI staining, we revealed that *ex-LacZ* was elevated in the primary pigment cells, compared to all other cell types (photoreceptor, cone, and interommatidial cells) (**Fig. 1A-K**). This was confirmed using two independent Yki/Sd transcription reporters; *bantam-LacZ (ban-lacZ)* and *DIAP1-GFP^4.3^*, both of which were substantially higher in primary pigment cells compared to other eye cells (**Fig. 1L-O and S1**). Using the *DIAP1-GFP^4.3^* reporter, we revealed that the asymmetry in Yki/Sd activity increases throughout pupal eye development from 19h APF to 42hAPF (**Fig. S1**).

### Scalloped is required for basal transcription in primary pigment cells of the *Drosophila* eye

Next, we assessed the genetic requirement of both Yki and Sd for transcription in pupal primary pigment cells. *yki* mutant eye cells grow poorly and are eliminated by apoptosis [52], and therefore primary pigment cells lacking *yki* were unable to be studied in the pupal eye. To overcome this limitation, we recombined a *yki* null allele with a mutation in the *Death-associated APAF1-related killer* (*dark)* gene, a pro-apoptotic regulator, whose mutation prevents cell death [53, 54]. As expected, expression of *ex-LacZ* and *DIAP1-GFP^4.3^* were unaltered in control *dark* single mutant clones (**Fig. 2A,E, and S2B,E)**. However, *ex-LacZ, ban-lacZ* and *DIAP1-GFP^4.3^* were all undetectable in *yki*, *dark* double mutant pupal eye clones 24h APF (**Fig. 2C-G, S2A and S2D**). Next, we examined a role for Yki’s DNA-binding partner Sd in the regulation of *ex* and *ban*. In larval eye discs, Sd is dispensable for tissue growth and transcription of target genes in the disc proper, while it is required for cell viability and gene expression in the peripodial epithelium [48, 55]. Interestingly, *ex-LacZ, ban-LacZ* and *DIAP1-GFP^4.3^* were all reduced in *sd* mutant primary pigment cells of the pupal eye (**Fig. 2H-K, S2C and S2F)**. In contrast to *yki* clones, where target gene transcription was undetectable, *sd* mutant primary pigment cells displayed residual transcription of each reporter gene (approximately 25-50% of that observed in wild-type cells). This result was confirmed by RNAi-mediated depletion of Sd, which also caused a partial reduction in *DIAP1-GFP^4.3^* abundance (**Fig. S2G-J**). Collectively, these experiments indicate that Sd is required to promote robust transcription of classical Yki/Sd target genes in primary pigment cells but is not essential for their expression. Further, this data suggests that in *yki* mutant clones, Sd’s well-defined default transcription repressor function dominates to limit Yki/Sd target gene expression [55, 56].

**Figure 2.**
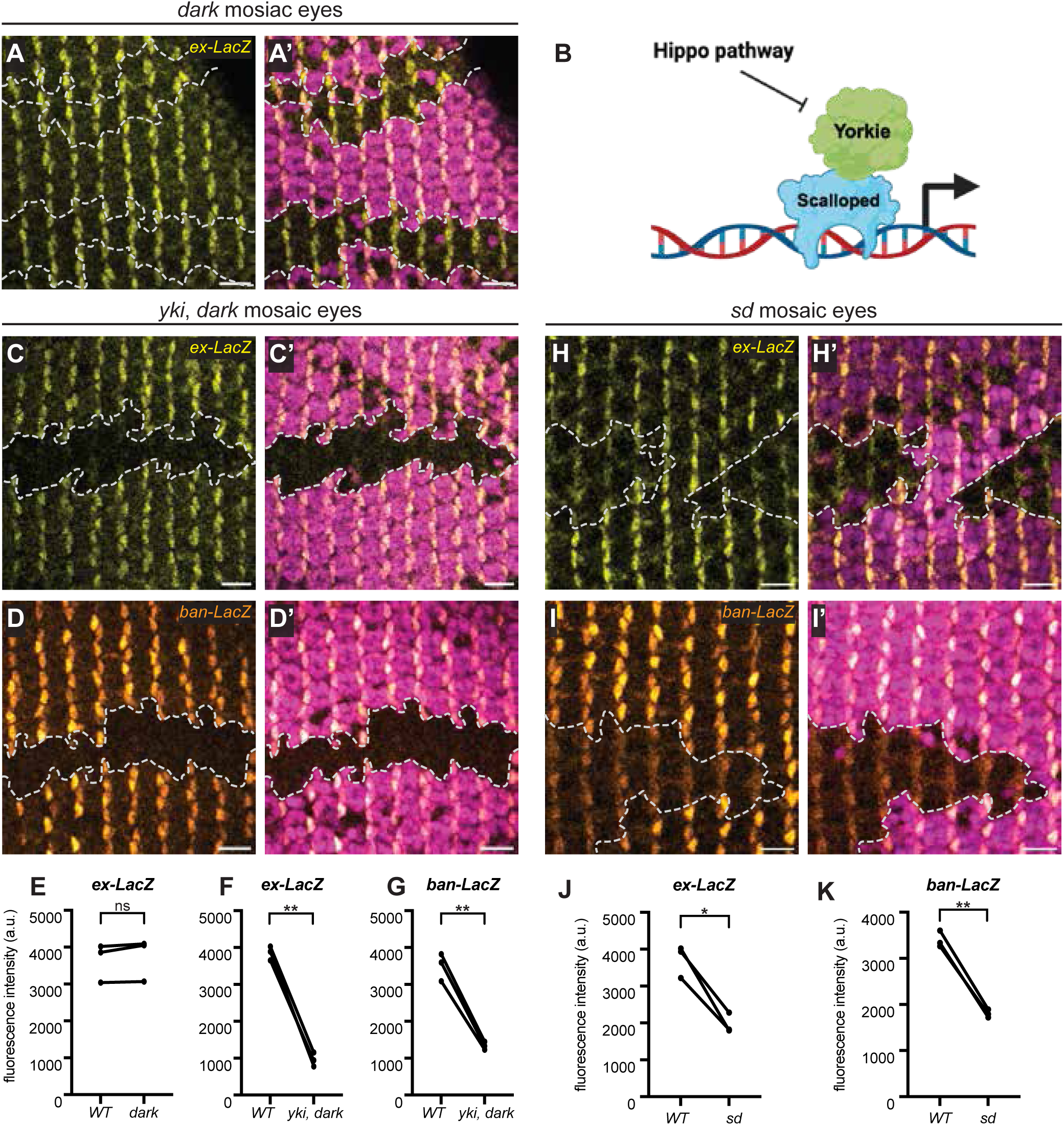
Scalloped is required for basal transcription in primary pigment cells of the *Drosophila* eye. (**A, C, D, H and I**) Pupal *Drosophila* eyes 24h APF harbouring patches of mutant tissue (RFP negative) for the following genes: *dark* (A), yki, *dark* (C and D), *sd* (H and I). Homozygous wildtype or heterozygous tissue (RFP positive) is in pink in the merged images, *ex-LacZ* is in yellow (A, C and H) and *ban-LacZ* is in orange (D and I). Clones are outlined by dashed white lines, scale bars = 10μm. **(B)** Schematic diagram of the downstream Hippo pathway transcription effectors, Yki and Sd. (**E, F, G, J and K**) Charts showing quantification of *ex-LacZ* (E, F and J) and *ban-LacZ* (G and K) fluorescence intensity in wildtype and mutant primary pigment cells within the same eye discs. n=3 for each sample, * indicates P-value<0.05, ** indicates P-value<0.005 and ns indicates P-value>0.05, assessed by paired t-tests.

### Yorkie and Scalloped are dispensable for cell fate in the pupal eye

Given that Yki/Sd activity is elevated in primary pigment cells and Yki/Sd are required for robust target gene expression in these cells, we assessed whether they are required for primary pigment cell fate. To do this, we assessed expression of two key markers of the primary pigment cell fate, the transcription factors DPax2 [10] and BarH1 [57, 58]. However, both proteins were expressed at normal levels in *yki, dark* clones (**Fig. 3A,B**). Similarly, Sd was dispensable for primary pigment cell fate, as BarH1 and DPax2 were expressed at normal levels and primary pigment cell shape (**Fig. S3A,B**). As a further indication that primary pigment cell fate was unaffected by loss of Yki/Sd activity, the distinctive “kidney” shape of primary pigment cells was unperturbed in both *yki*, *dark* clones and *sd* clones (**Fig. S3E,F**). In the same experiments, we examined the fate of cone cells and bristle cells by assessing Cut and dPax2 expression and found that both *yki* and *sd* were also dispensable for the fate of these cell types (**Fig. 3C,D and S3C,D,G, H**).

**Figure 3.**
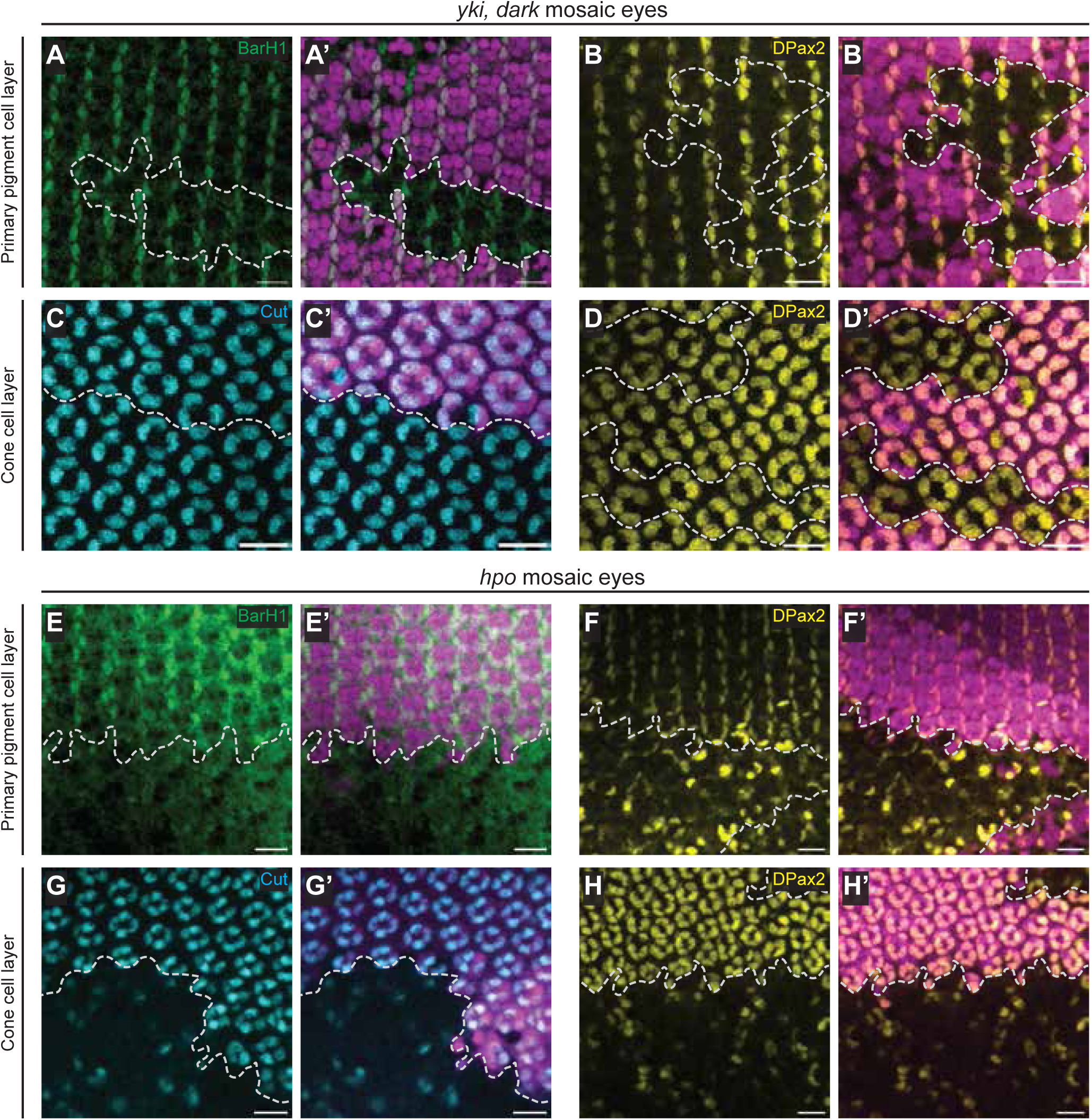
Hippo signalling promotes the cone and primary pigment cell fates of the *Drosophila* eye. (**A-H**) Confocal microscope images of *Drosophila* pupal eyes 24h APF at the primary pigment cell nuclei layer (A, B, E and F), and the cone cell nuclei layer (C, D, G and H). The tissues harbour patches of mutant tissue (RFP negative) for either *yki*, *dark* (A-D), or *hpo* (E-H), while homozygous wildtype or heterozygous tissue (RFP positive) is in pink in the merged images. BarHI is in green (A and E), DPax2 is in yellow (B, D, F and H) and Cut is in cyan (C and G). Clones are outlined by dashed white lines, scale bars = 10μm.

### Hippo signalling promotes the cone and primary pigment cell fates of the *Drosophila* eye

Previously, the Hippo pathway was found to be dispensable for photoreceptor cell fate in the third instar larval eye, as differentiation of these cells occurred normally in tissue lacking core Hippo pathway proteins (*hpo* and *sav*), where Yki/Sd transcription is elevated [17, 18, 21, 23, 24]. To examine a potential role for Hippo signalling at later stages of eye development, we assessed cell fate in *hpo* mutant pupal eye clones. Initially, we evaluated cone and primary pigment cells by assessing expression of the Cut and BarHI transcription factors, respectively. Strikingly, both Cut-positive cone cells and BarHI-positive primary pigment cells were almost completely absent from *hpo* eye clones 24h APF (**Fig. 3E and G**). This did not reflect a delay in differentiation as the same phenotype was also observed later in eye development (40h APF) (**Fig. S4A,C**). These phenotypes were not caused by a loss of these cells from the epithelium, as DAPI-staining revealed cell nuclei in both the cone cell and primary pigment cell layers of *hpo* mutant clones (**Fig. S4A-D**), and is consistent with the fact that *hpo* eye cells are resistant to programmed cell death [17, 18, 21, 23–25].

To investigate this further, we assessed expression of *dPax2*, which acts upstream of both Cut and BarH1 in cone and primary pigment cells, respectively [10]. DPax2 protein was almost completely absent from the cone cell layer and primary pigment cell layers of *hpo* mutant clones 24h APF (**Fig. 3F and H).** We did detect some DPax2-positive cells in *hpo* mutant pupal eye clones, especially in the primary pigment cell layer (**Fig. S4B,D**). Given the circular shape of the nuclei of these DPax2-positive cells, they are likely to be bristle cells [59], as opposed to primary pigment cells, whose nuclei have a slightly elongated oval shape. Indeed, bristle cells also express DPax2 and this transcription factor is essential for bristle cell development [60]. To determine whether DPax2’s absence from *hpo* mutant cone and primary pigment cells was transcriptional or post-transcriptional, we made use of a *dPax2* transcription reporter *Drosophila* strain. The *dPax2-LacZ* reporter was absent in both cone cells and primary pigment cells in *hpo* mutant pupal eye clones, indicating that *dPax2* is not transcribed in these cells (**Fig. 4A,B**).

**Figure 4.**
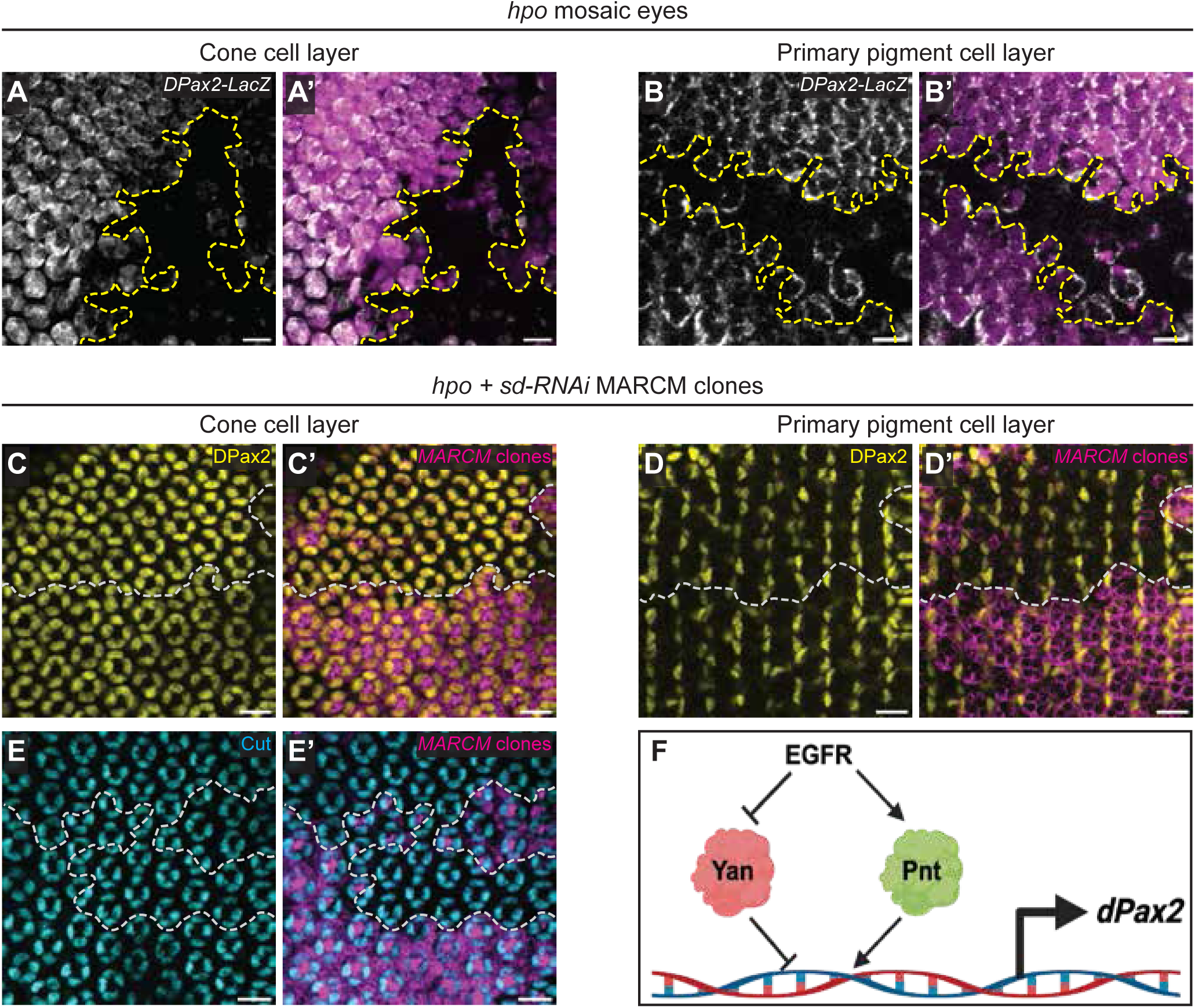
Yorkie/Scalloped hyperactivity antagonises cone and primary pigment cell fates of the *Drosophila* eye. (**A-E**) Confocal microscope images of *Drosophila* pupal eyes 24h APF at the cone cell nuclei layer (A, C and E) and the primary pigment cell nuclei layer (B and D). In (A and B), tissues harbour patches of *hpo* mutant tissue (RFP negative), while homozygous wildtype or heterozygous tissue (RFP positive) is in pink in the merged images. In (C-E), tissues harbour MARCM clones that are *hpo* mutant and express Sd RNAi (RFP positive – pink in the merged images), while control tissue does not express RFP. *DPax2-LacZ* is in greyscale (A and B) DPax2 is in yellow (C and D) and Cut is in cyan (E). Clones are outlined by dashed white lines, scale bars = 10μm. (**F**) Schematic diagram of the downstream Notch and EGFR pathway transcription effectors, Su(H), Yan, Pnt and Lz, which regulate *DPax2* transcription.

### Hippo signalling is dispensable for the induction of cone and primary pigment cells

The differentiation defects observed in *hpo* mutant eyes 24h APF could be caused either by a lack of cell fate induction or a lack of cell fate maintenance, given that cone cells first arise during third instar larval development [61]. To assess this, we examined the impact of *hpo* loss on cell fates in third instar larval eye imaginal discs. Cells expressing both the cone cell marker Cut and its upstream transcription factor DPax2 were readily observable in the posterior regions of *hpo* mutant larval eye imaginal disc tissue (**Fig. S5A,B**). In addition, *dPax2-LacZ,* was active in *hpo mutant* larval eye clones (**Fig. S5C**). This indicates that Hippo signalling is not required for the initial induction of cone cell fate in the larval eye but is required to maintain cone cell fate later in development.

Next, we investigated whether Hippo signalling is required for appropriate differentiation of additional *Drosophila* eye cell fates. *hpo* mutant pupal eyes are known to possess substantially more interommatidial cells, which we confirmed by detecting E-cad to mark cell outlines, and the transcription factor Lozenge (Lz), which marks pigment cells (**Fig. S6A,B**). We also assessed bristle cells, by detecting Cut expression and photoreceptor cells, by detecting Embryonic lethal abnormal vision (ELAV) expression. Bristle cell fate was unaffected, as Cut-positive bristle cells were still readily detectable in *hpo* eye clones 24h APF, even if the pattern of these cells was affected (**Fig. S6C**). Similarly, photoreceptor cell fate was largely unaffected in *hpo* eye clones 24h APF, as determined by detection of ELAV, although the distribution of these cells was perturbed, presumably at least partly because of the increased interommatidial cells. This result is consistent with previous analysis of photoreceptor cells in *wts* mutant pupal eyes [47]. Collectively, these data indicate that appropriate Hippo signalling is required for the maintenance of both the cone and primary pigment cell fates, but not other eye cell fates.

### Yorkie/Scalloped hyperactivity antagonises cone and primary pigment cell fates

Defective Hippo signalling causes nuclear accumulation of Yki, which in turn hyperactivates Sd-dependent transcription. Accordingly, phenotypes that result from Hippo signalling perturbation, like eye overgrowth, are completely suppressed by *sd* deficiency [38–40]. To determine whether the failure to maintain cone and primary pigment cell fate defects in *hpo* mutant eyes, was caused by Yki/Sd hyperactivity, we depleted Sd in *hpo* clones using the MARCM technique [62] and RNA interference [63, 64]. RNAi-mediated depletion of Sd in *hpo* mutant clones completely restored the expression of Cut and DPax2 in cone cells, as well as DPax2 in primary pigment cells (**Fig. 4C-E**). Furthermore, the ordered array of these cell types was largely restored in *hpo* mutant clones expressing Sd RNAi. Therefore, Hippo signalling promotes the maintenance of both cone and primary pigment cell fates in the pupal eye by repressing its canonical downstream transcription regulatory proteins, Yki and Sd.

### Hippo signalling promotes cone and primary pigment cell fate by repressing Yki-Sd-regulated transcription of *yan*

*dPax2* transcription is regulated by multiple signalling pathways and their cognate transcription factors: Notch signalling activates *dPax2* transcription via Su(H), while EGFR signalling promotes *dPax2* transcription by activating Pnt and inhibiting the Yan transcriptional repressor [11, 13]. In addition, Lz can activate *dPax2* transcription [11, 16] (**Fig. 4F**). As such, we considered the possibility that Yki/Sd might regulate the expression of one or more of these factors, which could explain the cell fate defects in *hpo* pupal eyes. We already established that Lz was expressed in *hpo* pupal eye cells at similar levels as wildtype cells (**Fig. S6B**). Therefore, we searched genes that we previously identified as Yki/Sd target genes in larval *Drosophila* eyes, using targeted DamID-seq [65]. Both Yki-Dam and Sd-Dam were significantly enriched at the *yan* locus compared to the Dam alone control, consistent with the notion that *yan* is a Yki/Sd target gene (**Fig. 5A**). Given that Yan overexpression can repress *dPax2* expression [11], we assessed its abundance in both *wts* and *hpo* pupal eye clones 24h APF and found that Yan protein abundance was elevated compared to wildtype cells in both experiments (**Fig. 5A,C)**. This effect was restricted to the pupal eye, as Yan abundance was not elevated in *hpo* mutant third instar larval eye clones (**Fig. S5D**). Elevation of Yan in Yki hyperactive pupal eye tissues resulted from increased *yan* transcription as a *yan-LacZ* transcription reporter [66] was elevated in *wts* pupal eye clones (**Fig. 5B**). Finally, RNA-mediated interference of Sd in *hpo* mutant clones reverted the elevated Yan expression back to baseline levels, indicating that Yki/Sd drive elevated *yan* transcription in pupal eye tissues with defective Hippo signalling (**Fig. 5D**).

**Figure 5.**
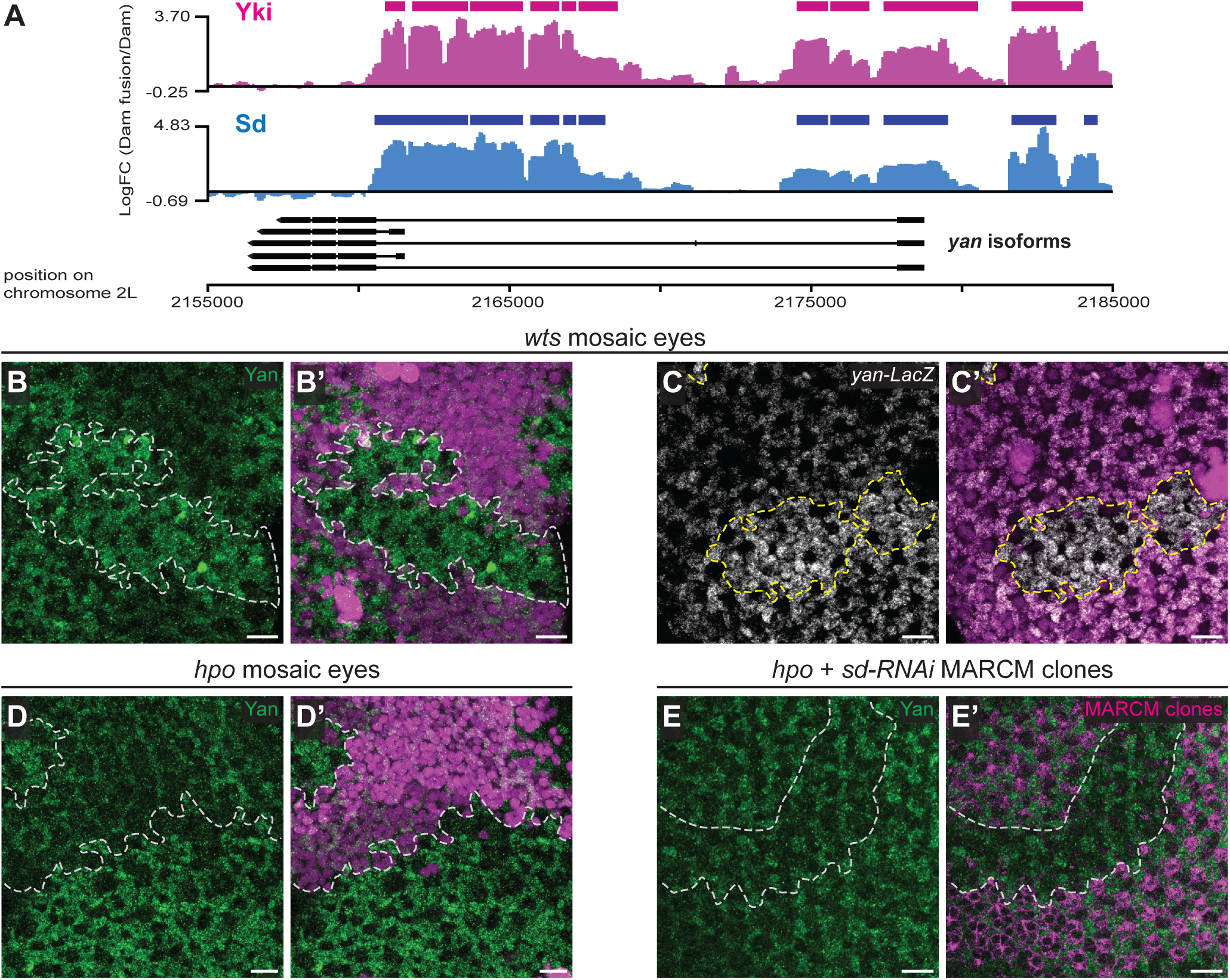
Hippo signalling limits expression of the Yan transcription repressor in the pupal *Drosophila* eye. (**A**) Genome-binding profiles of the *yan* gene locus by Yki and Sd, as determined by targeted DamID-seq. Yki binding peaks are in pink and Sd binding peaks are in blue. The enrichment of Yki-Dam and Sd-Dam normalised to the Dam-alone control is shown on the y-axis. Solid bars represent peaks where methylation by Yki-Dam or Sd-Dam scored as significant. (**B-E**) Confocal microscope images of *Drosophila* pupal eyes 24h APF. The tissues harbour patches of mutant tissue (RFP negative) for either *wts* (B and C), or *hpo* (D), while homozygous wildtype or heterozygous tissue (RFP positive) is in pink in the merged images. In (E), tissues harbour MARCM clones that are *hpo* mutant and express Sd RNAi (RFP positive – pink in the merged image), while control tissue does not express RFP. Yan is in green (B, D and E) and *yan-LacZ* is in greyscale (C). Clones are outlined by dashed white or yellow lines, scale bars = 10μm.

Next, we tested whether ectopic Yan expression underpinned the loss of cone and primary pigment cells in *hpo* mutant pupal eyes. Strikingly, RNAi-mediated Yan depletion in *hpo* pupal eye clones largely restored BarHI expression, indicating the restoration of the primary pigment cell fate (**Fig. 6A**). Note, these clones did display irregular patterning of primary pigment cells, where the pairing of two primary pigment cells per ommatidium was often disrupted, as were the neat rows of primary pigment cell patterns. Additionally, Cut and DPax2 expression were both restored in *yan*-depleted, *hpo* mutant pupal eye clones, indicating the restoration of cone cells (**Fig. 6B,C**). Note that instead of a regular group of four cone cells per ommatidium, some *yan*-*RNAi* depleted *hpo* clones displayed more or less than four cone cells per ommatidium, suggestive of incomplete restoration of ommatidial patterning. Collectively, these experiments argue that Hippo signalling normally promotes cone and primary pigment cell fate in the pupal eye via a double repression mechanism, i.e. Hippo signalling represses Yki/Sd from transcribing *yan*, and in doing so prevents Yan from ectopically repressing the cone and primary pigment cell fates.

**Figure 6.**
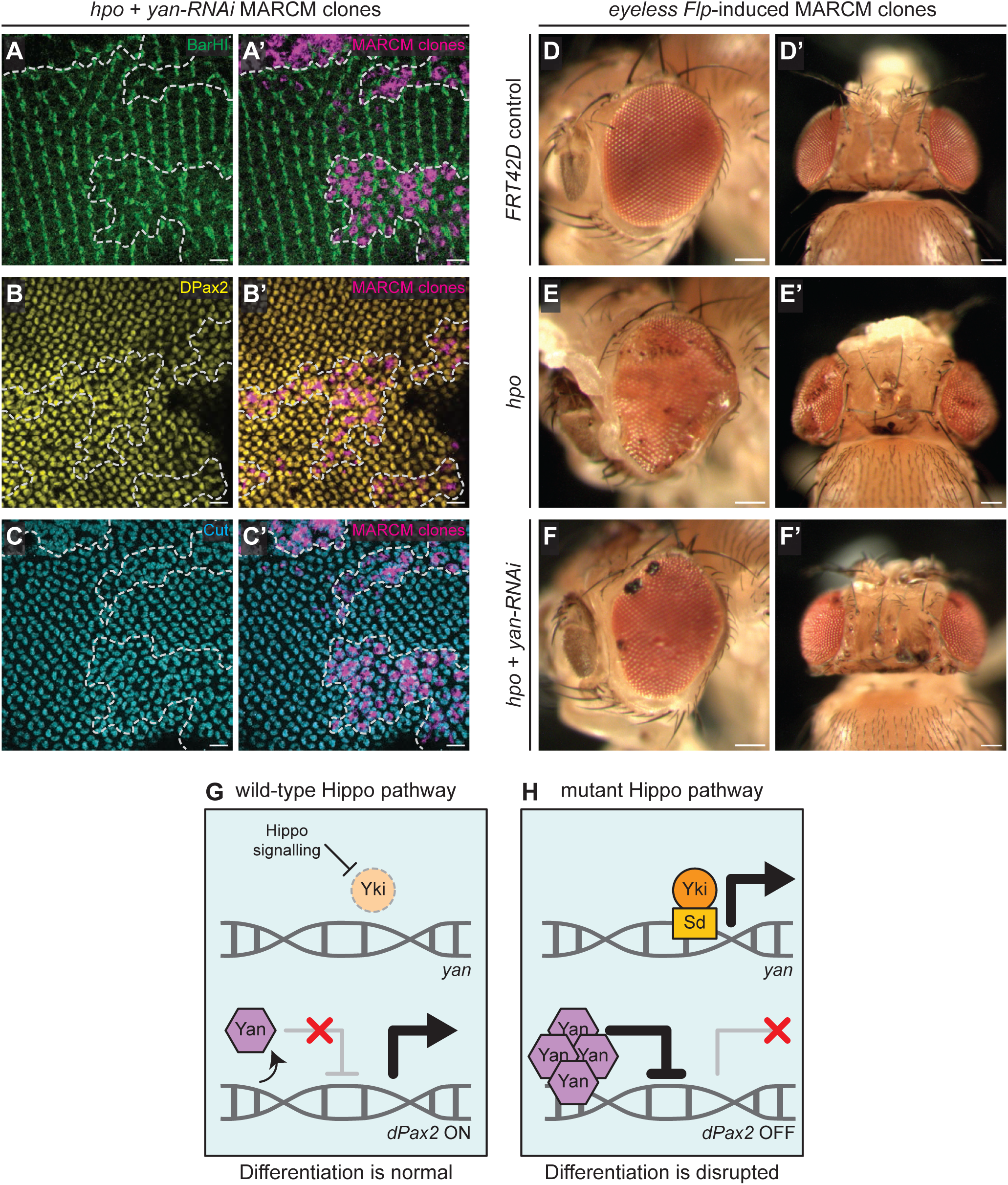
Hippo signalling promotes cone and primary pigment cell fate by repressing Yki-Sd-regulated transcription of *yan.* (**A-C**) Confocal microscope images of *Drosophila* pupal eyes 24h APF at the primary pigment cell nuclei layer (A) or the cone cell nuclei layer (B and C). The tissues harbour MARCM clones that are *hpo* mutant and express Yan RNAi (RFP positive – pink in the merged images), while control tissue does not express RFP. BarH1 is in green (A), DPax2 is in yellow (B) and Cut is in cyan (C). Clones are outlined by dashed white lines, scale bars = 10μm. (**D-F**) Adult *Drosophila* that harbour eyFlp-generated MARCM clones; either control (D), *hpo* mutant (E) or *hpo* mutant that express Yan RNAi (F). **(G and H)** Models of how Hippo signalling and Yan impact DPax2 expression and cell differentiation in the *Drosophila* eye.

Finally, given that Yan depletion reversed cell fate defects caused by Hippo pathway perturbation to a similar extent as Sd depletion, we examined whether Yan depletion could impact tissue overgrowth caused by disrupted Hippo signalling. *hpo* loss causes strong overgrowth of all imaginal disc tissues, which is visible in adults tissues such as the eye, head and thorax [67]. *hpo* mosaic eyes generated by the *eyeless-Flp* MARCM technique [68], were substantially overgrown compared to controls, as was the head cuticle (**Fig. 6D,E**). RNA-mediated depletion of Yan in *hpo* clones generated via the *eyeless-Flp* MARCM technique strongly suppressed eye and head overgrowth (**Fig. 6F**). Adult eyes harbouring Yan RNAi, *hpo* clones exhibited dark patches, which were presumably caused by Yan depletion, given that this was observable in Yan RNAi clones, but not *hpo* clones (**Fig. 6E and S7A**), and similar phenotypes have been reported previously for *yan* mutant adult eye clones [69]. The impact of Yan depletion on *hpo* clone overgrowth was restricted to the eye and head cuticle, as adult thorax Yan RNAi, *hpo* clones generated via the *hs-Flp* MARCM technique still strongly overgrew and resembled published reports of *hpo* mutant thorax clones [21, 67] (**Fig. S7B**). Yan depletion in these experiments was confirmed by immunostaining larval eye discs (**Fig. S7C**). Collectively, this shows that Yan depletion reverts both cell fate defects and overgrowth caused by *hpo* mutations in the *Drosophila* eye but not does impact Yki/Sd-driven overgrowth of non-eye tissues.

## DISCUSSION

By studying Hippo signalling activity in the developing *Drosophila* eye, we found that pathway activity is patterned in the pupal post-mitotic phase of development, despite having no major activity patterns during the larval growth phase of development. Yki-Sd activity was strongly elevated in primary pigment cells relative to all other eye cells, i.e. secondary and tertiary pigment cells, bristles, photoreceptors and cone cells. This implies that Hippo signalling activity is lower in primary pigment cells, although this requires formal evaluation with reagents that are currently not available, e.g. in vivo activity reporters of the kinases Hpo and/or Wts. It is also currently unclear whether elevated Yki/Sd activity in primary pigment cells has a functional role. Pupal eye cells, including primary pigment cells lacking *yki*, were unable to be recovered indicating that Yki is required for their survival. However, when we generated *yki* mutant primary pigment cells that were impervious to cell death we observed no obvious impacts to cell fate. Nevertheless, we cannot rule out the possibility that Yki/Sd drive a gene expression program that contributes to primary pigment cell function without obviously altering the fate of these cells.

What is apparent from our studies is that tight control of Yki/Sd activity by the Hippo pathway is essential in the post-mitotic phase of eye development, as Yki-Sd hyperactivity caused a profound loss of fate of both primary pigment cells and cone cells. The role of Hippo signalling in cone cell fate was limited to cone cell fate maintenance, as opposed to induction, as these cells were still induced normally in eye tissue lacking *hpo* during larval development. Therefore, Hippo signalling is required to actively maintain, but not induce, cone cells, suggesting that the maintenance of the fate of these cells is an active process. Interestingly, Hippo signalling is also required to maintain the fate of “yellow” R8 photoreceptor cells in the *D. melanogaster* eye, which express the light sensitive protein Rhodopsin 6 which is sensitive to green light [70]. Furthermore, Hippo signalling is required to maintain the fate of photoreceptor cells that have perturbed *Rbf* function [71]. Therefore, while the majority of studies have focused on the role of Hippo signalling in controlling the growth of many tissues from diverse species, it might also have a broader than appreciated role in cell fate maintenance.

At the mechanistic level, we found that Yki-Sd hyperactivity induces cone and primary pigment cell fate loss by driving elevated expression of the Yan transcription repressor, an essential regulator of transcription downstream of the EGFR pathway. Targeted DamID-seq studies revealed that Yki and Sd each to the *yan* gene, while both a *yan* transcription reporter and Yan protein were elevated in *hpo* mutant pupal eye clones in a Sd-dependent fashion. Furthermore, Yan depletion in *hpo* mutant pupal eye cells largely restored the fate of both cone and primary pigment cells. This highlights the importance of maintaining Yan levels and transcription control by the EGFR pathway for cell fate maintenance. Furthermore, the increase in interommatidial cells at the expense of cone and primary pigment cells in *hpo* eyes is consistent with the model where interommatidial cell fate is the “default” fate for a common pool of precursor cells [3, 72, 73].

Yan is an ETS family transcriptional repressor that competes for binding to common genomic loci with another ETS family transcription factor activator Pnt, to regulate transcription of genes that promote differentiation of photoreceptor cells [66, 74, 75]. Yan and Pnt also regulate the cone cell fate by binding to the enhancer region of *dPax2* at multiple sites and regulating its transcription [11, 13]. EGFR signalling both downregulates Yan and upregulates Pnt to alter the cellular ratio of Yan/Pnt, which is essential for dictating whether their shared target genes are either activated or repressed [76]. Our experiments suggest that in *hpo* mutant pupal eyes, ectopic elevation of Yan by Yki-Sd tips the Yan/Pnt ratio in favour of Yan, which represses *dPax2* transcription, and causes cone cell de-differentiation.

Our study provides yet another example of the close functional relationship that exists between the Hippo and EGFR pathways. Multiple reports in both *Drosophila* and mammals have shown that these pathways regulate the expression of a shared transcriptome and cooperatively influence cell proliferation and organ growth, and cancer therapy resistance [77–79]. Consistent with this, multiple recent oncology studies have revealed enhanced anti-tumour efficacy when EGFR pathway and Hippo pathway targeted therapies are combined [80–84]. Here, we show that the close functional relationship between the EGFR and Hippo pathways is also essential for the control of cell fate during development.

## MATERIALS AND METHODS

### D. melanogaster strains

The following *D. melanogaster* stocks were used, some of which were from the Bloomington Drosophila Stock Centre (BDSC): *sd ^47M^ FRT19A* [85]; *FRT42D yki ^B5^* [52]; *FRT42D dark ^22B1^* [54]; *FRT42D hpo ^5.1^* [86]; *FRT82B wts ^X1^* [20]; *Ubi-RFP.NLS, hs-FLP, FRT19A* (BDSC, # 31418); *arm-LacZ, FRT19A* [87]; *FRT42D ubi-RFP.NLS* (BDSC, #35496); *FRT 42D arm-LacZ* (BDSC, #7372); *FRT82B ubi-RFP.NLS* (BDSC, #30555); *FRT42D tubP-GAL80* (BDSC, #9917); *ey-FLP.N (X)* (BDSC, #5580); *hs-FLP.D5 (X)* (BDSC, #55815); *hs-FLP.D5 (III)* (BDSC, #55813); *ex-lacZ* (BDSC, #44248); *ban-LacZ* (BDSC, #10154); *dPax2-LacZ* (BDSC, #33840); *yan-LacZ* (BDSC, 11359); *DIAP1-GFP ^4.3^* [40]; *De-GAL4 (III*) (BDSC, #29650); *tubP-GAL4 (III)* (BDSC, #5138); *UAS-mCD8-mRFP (III)* (BDSC, #27399); *UAS-yan-*RNAi (BDSC, #34909); *UAS-sd-RNAi ^(N+C)^* [40]. The following lines were generated by meiotic recombination: *FRT42D yki ^B5^ dark ^22B1^*; *ex-LacZ FRT42D ubi-RFP.NLS; DE-GAL4 UAS-mCD8-mRFP (III)*; and *tubP-GAL4 UAS-mCD8-mRFP (III)*.

### *D. melanogaster* husbandry, heat shock and staging

*Drosophila melanogaster* were raised at 25°C on a standard diet made with semolina, glucose (dextrose), raw sugar, yeast, potassium tartrate, nipagen (tegosept), agar, propionic acid and calcium chloride. To ensure that nutritional availability was not limiting, all animals were fed in excess food availability, while carefully controlling for crowding in the vials. Only female flies were used in experiments using the *sd^47M^* allele and experiments that investigated effects in the adult eye. For all other experiments, both male and female flies were used. To generate homozygous mutant clones in otherwise heterozygous whole organisms, the FLP-FRT system was used [68, 88, 89], using either *eyeless-FLP* (*ey-FLP*) or *heatshock-FLP* (*hs-FLP*). For experiments using *hs-FLP*, the heat shock protocols used were dependent on the mutant allele(s) being investigated, as follows: *dark^22B1^, hpo^5.1^, sd^47M^,* and *yki^B5^ dark ^22B1^ -* heat shocked at 37°C 48h after egg laying for 15 minutes; *wts^X1^* - heat shocked at 37°C 72h after egg laying for 10 minutes. To generate MARCM clones in otherwise heterozygous organisms [62, 90], both *ey-FLP* and *hs-FLP* were used (heat shocked at 37°C 48h after egg laying for 15 minutes). Wandering third instar larvae were used for all larval dissections. For pupal dissections, white pre-pupae were isolated and incubated at 25 C° for the desired time before they were dissected.

### Immunostaining and confocal microscopy

All tissues were dissected in PBS and put immediately on ice. Fixation and immunofluorescence for larval tissues were performed as described in [91, 92]. For pupal tissues, the PAXD buffer was used during immunostaining, as described in [93]. All tissues were fixed in 4% Paraformaldehyde, permeabilised and incubated with antibodies according to the reagents Table. Tissues were mounted in Vectashield antifade medium and confocal microscope images were taken on an Olympus FV3000 microscope, using either a 30x silicon objective or a 60x oil objective. To image adult *D. melanogaster*, 1 week-old adults were flash frozen on dry ice, mounted on wet tissue and imaged with an Infinity 1 camera at 4x magnification.

### Adult *D. melanogaster* microscopy

Adult *D. melanogaster* were frozen at -20°C overnight and imaged using a Zeiss Stemi 305 with Axiocam at 4X zoom as in [94, 95].

### Image analysis and statistics

Fluorescence microscopy images were analyzed using FIJI [96] and all statistical analyses were conducted in GraphPad Prism. To quantify fluorescence intensity, Regions of Interest (ROI) were drawn for cells of interest (eg. Primary Pigment cell) using FIJI, and each cell’s genotype was assigned ‘WT’ or ‘mutant’ according to the corresponding fluorescent marker. An average fluorescence value was recorded for WT cells and mutant cells within the same tissue. For comparisons between two genotypes, t-tests were conducted using paired analyses, with the WT cells serving as internal controls. For comparisons between more than two cell types one-way ANOVA tests were performed with Tukey-Kramer tests to correct for multiple comparisons. The significance value was set at 0.05, and a P-value of 0.05 or lower was considered statistically significant. Images were cropped and rotated in FIJI, while dashed lines demarcating WT-mutant borders were drawn in Adobe Illustrator.

## Reagents Table

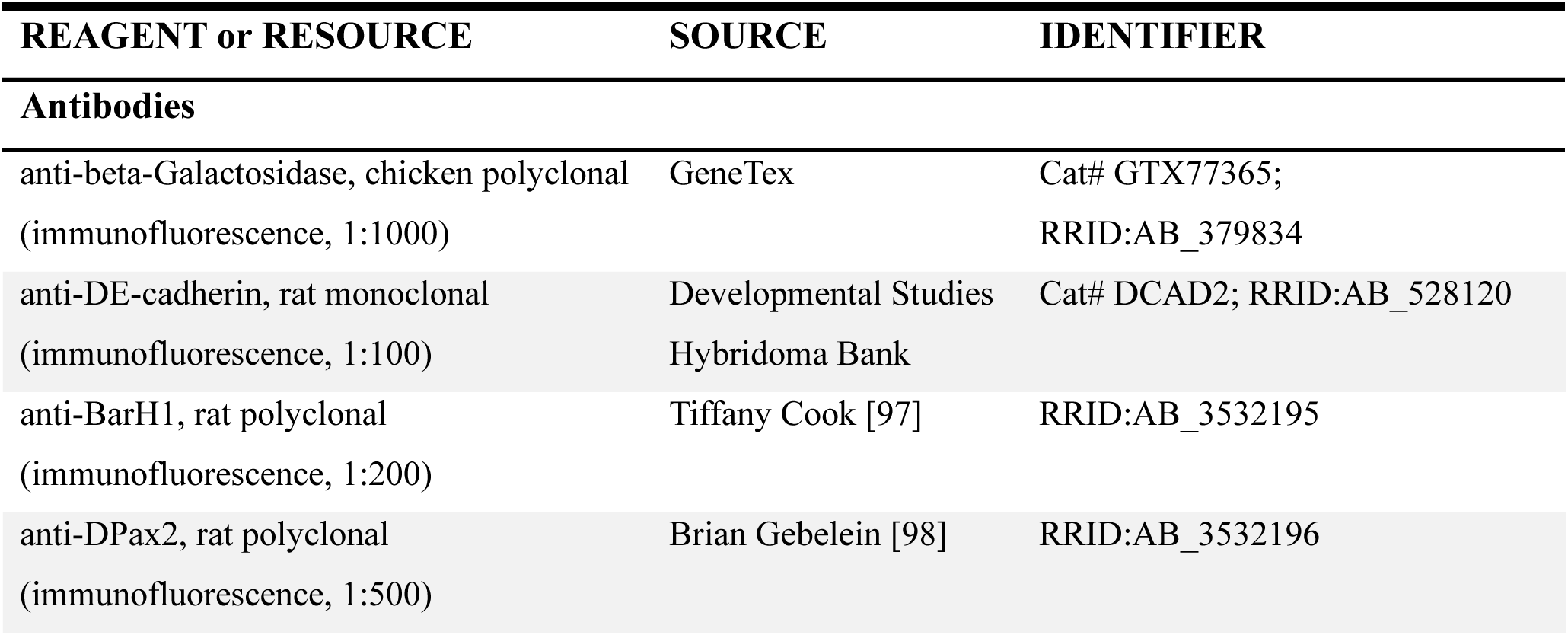
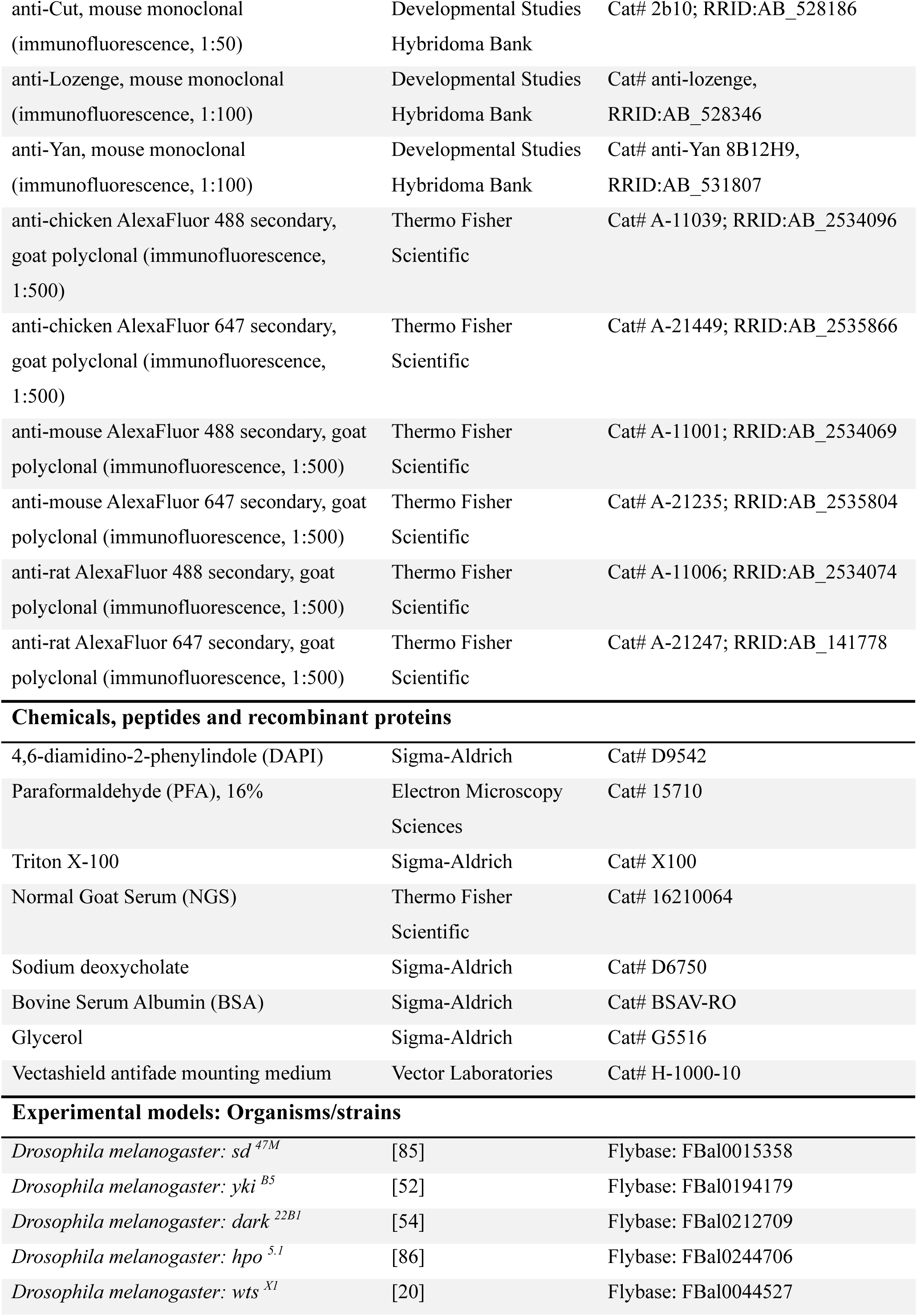
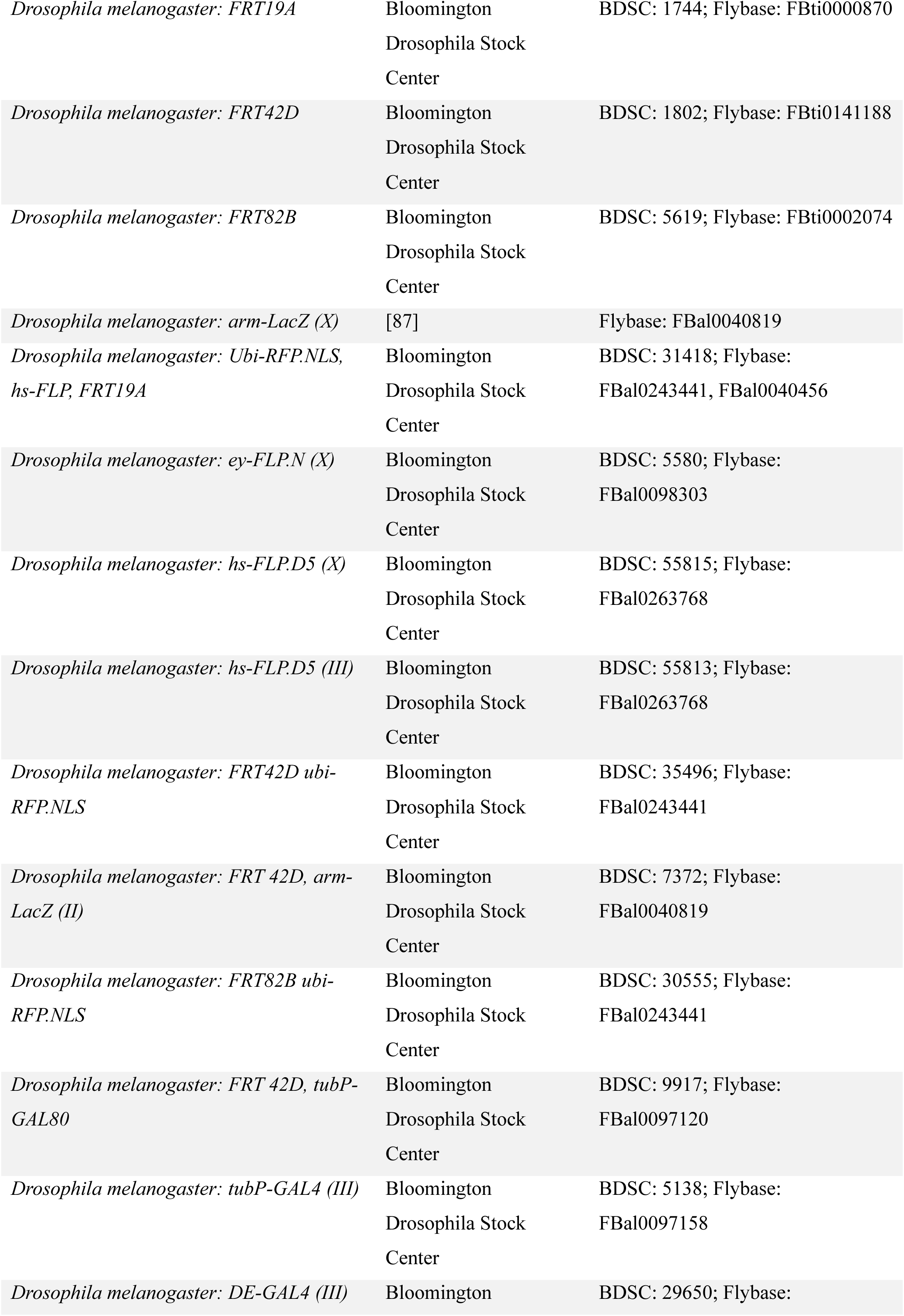
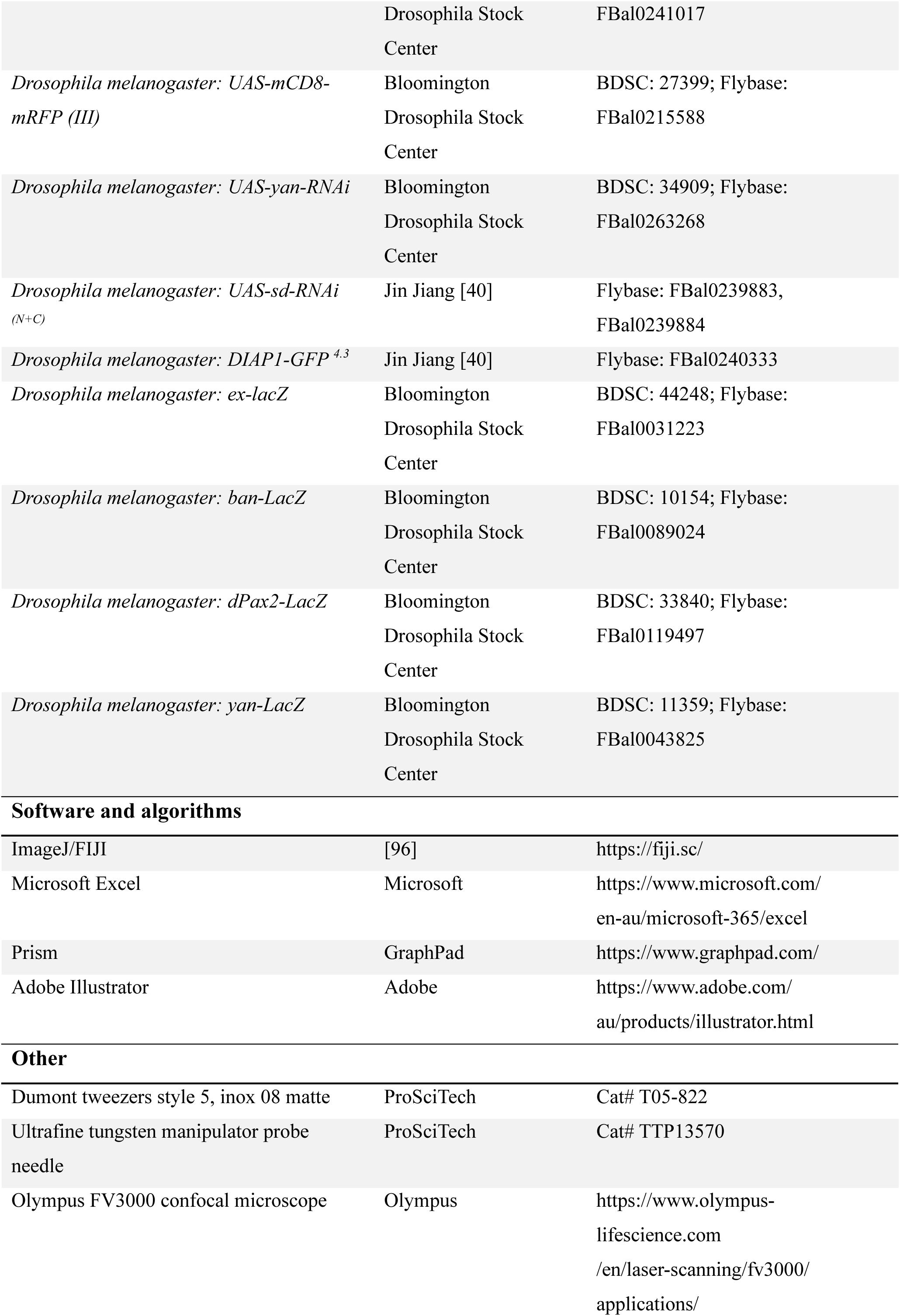
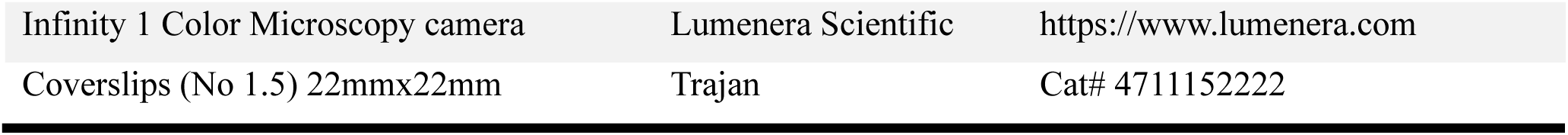

## ACKNOWLEDGEMENTS

We thank members of the Harvey lab for discussions, and T. Cook, B. Gebelein, J. Jiang, D. Pan, N. Tapon, T. Xu, the Bloomington *Drosophila* Stock Center, the Vienna *Drosophila* RNAi Center, the Australian *Drosophila* Research Support Facility, and the Developmental Studies Hybridoma Bank for *D. melanogaster* stocks and antibodies. K.F.H was supported by a Senior Research Fellowship (APP1078220) and Investigator grant (APP1194467) from the National Health and Medical Research Council (NHMRC) and A.J.S.H. was supported by a Peter Mac Postgraduate Award. This research was supported by the Australian Research Council (DP180102044) and the NHMRC (APP1157737). We acknowledge the Peter Mac Centre for Advanced Histology and Microscopy and Research Laboratory Support Services and support to them from the Peter MacCallum Cancer Foundation and the Australian Cancer Research Foundation.

## REFERENCES

1. Ready DF, Hanson TE, Benzer S. Development of the Drosophila retina, a neurocrystalline lattice. Dev Biol. 1976;53(2):217–40. doi: 10.1016/0012-1606(76)90225-6. PubMed PMID: 825400.

2. Charlton-Perkins MA, Friedrich M, Cook TA. Semper’s cells in the insect compound eye: Insights into ocular form and function. Dev Biol. 2021;479:126–38. Epub 20210731. doi: 10.1016/j.ydbio.2021.07.015. PubMed PMID: 34343526; PubMed Central PMCID: PMCPMC8410683.

3. Kumar JP. Building an ommatidium one cell at a time. Dev Dyn. 2012;241(1):136–49. doi: 10.1002/dvdy.23707. PubMed PMID: 22174084; PubMed Central PMCID: PMCPMC3427658.

4. Waddington CH, Perry MM. The ultra-structure of the developing eye of Drosophila. Proceedings of the Royal Society of London Series B Biological Sciences. 1960;153(951):155–78. doi: 10.1098/rspb.1960.0094.

5. Cagan RL, Ready DF. The emergence of order in the Drosophila pupal retina. Dev Biol. 1989;136(2):346–62. doi: 10.1016/0012-1606(89)90261-3. PubMed PMID: 2511048.

6. Tomlinson A, Ready DF. Neuronal differentiation in Drosophila ommatidium. Dev Biol. 1987;120(2):366–76. doi: 10.1016/0012-1606(87)90239-9. PubMed PMID: 17985475.

7. Freeman M. Reiterative use of the EGF receptor triggers differentiation of all cell types in the Drosophila eye. Cell. 1996;87(4):651–60. doi: 10.1016/s0092-8674(00)81385-9. PubMed PMID: 8929534.

8. Wolff T, Ready DF. The beginning of pattern formation in the Drosophila compound eye: the morphogenetic furrow and the second mitotic wave. Development. 1991;113(3):841–50. doi: 10.1242/dev.113.3.841. PubMed PMID: 1726564.

9. Carthew RW. Pattern formation in the Drosophila eye. Curr Opin Genet Dev. 2007;17(4):309–13. Epub 20070706. doi: 10.1016/j.gde.2007.05.001. PubMed PMID: 17618111; PubMed Central PMCID: PMCPMC2693403.

10. Fu W, Noll M. The Pax2 homolog sparkling is required for development of cone and pigment cells in the Drosophila eye. Genes Dev. 1997;11(16):2066–78. doi: 10.1101/gad.11.16.2066. PubMed PMID: 9284046; PubMed Central PMCID: PMCPMC316453.

11. Flores GV, Duan H, Yan H, Nagaraj R, Fu W, Zou Y, et al. Combinatorial signaling in the specification of unique cell fates. Cell. 2000;103(1):75–85. PubMed PMID: 11051549.

12. Pickup AT, Ming L, Lipshitz HD. Hindsight modulates Delta expression during Drosophila cone cell induction. Development. 2009;136(6):975–82. doi: 10.1242/dev.027318. PubMed PMID: 19234063.

13. Swanson CI, Evans NC, Barolo S. Structural rules and complex regulatory circuitry constrain expression of a Notch- and EGFR-regulated eye enhancer. Dev Cell. 2010;18(3):359–70. doi: 10.1016/j.devcel.2009.12.026. PubMed PMID: 20230745; PubMed Central PMCID: PMCPMC2847355.

14. Flores GV, Daga A, Kalhor HR, Banerjee U. Lozenge is expressed in pluripotent precursor cells and patterns multiple cell types in the Drosophila eye through the control of cell-specific transcription factors. Development. 1998;125(18):3681–7. Epub 1998/08/26. PubMed PMID: 9716533.

15. Yan H, Canon J, Banerjee U. A transcriptional chain linking eye specification to terminal determination of cone cells in the Drosophila eye. Dev Biol. 2003;263(2):323–9. doi: 10.1016/j.ydbio.2003.08.003. PubMed PMID: 14597205.

16. Canon J, Banerjee U. In vivo analysis of a developmental circuit for direct transcriptional activation and repression in the same cell by a Runx protein. Genes Dev. 2003;17(7):838–43. PubMed PMID: 12670867.

17. Tapon N, Harvey KF, Bell DW, Wahrer DCR, Schiripo TA, Haber DA, et al. salvador promotes both cell cycle exit and apoptosis in Drosophila and is mutated in human cancer cell lines. Cell. 2002;110(4):467–78. doi: 10.1016/S0092-8674(02)00824-3.

18. Pantalacci S, Tapon N, Leopold P. The Salvador partner Hippo promotes apoptosis and cell-cycle exit in Drosophila. Nat Cell Biol. 2003;5(10):921–7. PubMed PMID: 14502295.

19. Justice RW, Zilian O, Woods DF, Noll M, Bryant PJ. The Drosophila tumor suppressor gene warts encodes a homolog of human myotonic dystrophy kinase and is required for the control of cell shape and proliferation. Genes Dev. 1995;9(5):534–46. PubMed PMID: 7698644.

20. Xu T, Wang W, Zhang S, Stewart RA, Yu W. Identifying tumor suppressors in genetic mosaics: the Drosophila lats gene encodes a putative protein kinase. Development. 1995;121(4):1053–63. PubMed PMID: 7743921.

21. Wu S, Huang J, Dong J, Pan D. hippo encodes a Ste-20 family protein kinase that restricts cell proliferation and promotes apoptosis in conjunction with salvador and warts. Cell. 2003;114(4):445–56. PubMed PMID: 12941273.

22. Jia J, Zhang W, Wang B, Trinko R, Jiang J. The Drosophila Ste20 family kinase dMST functions as a tumor suppressor by restricting cell proliferation and promoting apoptosis. Genes Dev. 2003;17(20):2514–9. PubMed PMID: 14561774.

23. Kango-Singh M, Nolo R, Tao C, Verstreken P, Hiesinger PR, Bellen HJ, et al. Shar-pei mediates cell proliferation arrest during imaginal disc growth in Drosophila. Development. 2002;129(24):5719–30. PubMed PMID: 12421711.

24. Udan RS, Kango-Singh M, Nolo R, Tao C, Halder G. Hippo promotes proliferation arrest and apoptosis in the Salvador/Warts pathway. Nat Cell Biol. 2003;5(10):914–20. PubMed PMID: 14502294.

25. Harvey KF, Pfleger CM, Hariharan IK. The Drosophila Mst ortholog, hippo, restricts growth and cell proliferation and promotes apoptosis. Cell. 2003;114(4):457–67. PubMed PMID: 12941274.

26. Halder G, Dupont S, Piccolo S. Transduction of mechanical and cytoskeletal cues by YAP and TAZ. Nat Rev Mol Cell Biol. 2012;13(9):591–600. Epub 2012/08/17. doi: 10.1038/nrm3416. PubMed PMID: 22895435.

27. Zheng Y, Pan D. The Hippo Signaling Pathway in Development and Disease. Dev Cell. 2019;50(3):264–82. Epub 2019/08/07. doi: 10.1016/j.devcel.2019.06.003. PubMed PMID: 31386861.

28. Davis JR, Tapon N. Hippo signalling during development. Development. 2019;146(18). Epub 20190916. doi: 10.1242/dev.167106. PubMed PMID: 31527062; PubMed Central PMCID: PMCPMC7100553.

29. Pan Y, Alegot H, Rauskolb C, Irvine KD. The dynamics of Hippo signaling during Drosophila wing development. Development. 2018;145(20). Epub 2018/09/27. doi: 10.1242/dev.165712. PubMed PMID: 30254143; PubMed Central PMCID: PMCPMC6215397.

30. Enderle L, McNeill H. Hippo gains weight: added insights and complexity to pathway control. Science signaling. 2013;6(296):re7. Epub 2013/10/10. doi: 10.1126/scisignal.2004208. PubMed PMID: 24106343.

31. Harvey KF, Zhang X, Thomas DM. The Hippo pathway and human cancer. Nat Rev Cancer. 2013;13(4):246–57. Epub 2013/03/08. doi: 10.1038/nrc3458. PubMed PMID: 23467301.

32. Yu FX, Guan KL. The Hippo pathway: regulators and regulations. Genes Dev. 2013;27(4):355–71. Epub 2013/02/23. doi: 10.1101/gad.210773.112. PubMed PMID: 23431053; PubMed Central PMCID: PMC3589553.

33. Dong J, Feldmann G, Huang J, Wu S, Zhang N, Comerford SA, et al. Elucidation of a universal size-control mechanism in Drosophila and mammals. Cell. 2007;130(6):1120–33. PubMed PMID: 17889654.

34. Oh H, Irvine KD. In vivo regulation of Yorkie phosphorylation and localization. Development. 2008;135(6):1081–8. Epub 2008/02/08. doi: 10.1242/dev.015255. PubMed PMID: 18256197; PubMed Central PMCID: PMCPMC2387210.

35. Zhao B, Wei X, Li W, Udan RS, Yang Q, Kim J, et al. Inactivation of YAP oncoprotein by the Hippo pathway is involved in cell contact inhibition and tissue growth control. Genes Dev. 2007;21(21):2747–61. PubMed PMID: 17974916.

36. Oka T, Mazack V, Sudol M. Mst2 and Lats kinases regulate apoptotic function of Yes kinase-associated protein (YAP). J Biol Chem. 2008;283(41):27534–46. Epub 2008/07/22. doi: 10.1074/jbc.M804380200. PubMed PMID: 18640976.

37. Zhao B, Ye X, Yu J, Li L, Li W, Li S, et al. TEAD mediates YAP-dependent gene induction and growth control. Genes Dev. 2008;22(14):1962–71. PubMed PMID: 18579750.

38. Goulev Y, Fauny JD, Gonzalez-Marti B, Flagiello D, Silber J, Zider A. SCALLOPED interacts with YORKIE, the nuclear effector of the hippo tumor-suppressor pathway in Drosophila. Curr Biol. 2008;18(6):435–41. PubMed PMID: 18313299.

39. Wu S, Liu Y, Zheng Y, Dong J, Pan D. The TEAD/TEF family protein Scalloped mediates transcriptional output of the Hippo growth-regulatory pathway. Dev Cell. 2008;14(3):388–98. PubMed PMID: 18258486.

40. Zhang L, Ren F, Zhang Q, Chen Y, Wang B, Jiang J. The TEAD/TEF family of transcription factor Scalloped mediates Hippo signaling in organ size control. Dev Cell. 2008;14(3):377–87. PubMed PMID: 18258485.

41. Manning SA, Kroeger B, Deng Q, Brooks E, Fonseka Y, Hinde E, et al. The Drosophila Hippo pathway transcription factor Scalloped and its co-factors alter each other’s chromatin binding dynamics and transcription in vivo. Dev Cell. 2024;59(13):1640–54 e5. Epub 20240425. doi: 10.1016/j.devcel.2024.04.006. PubMed PMID: 38670104.

42. Kroeger B, Manning SA, Mohan V, Lou J, Sun G, Lamont S, et al. Hippo signaling regulates the nuclear behavior and DNA binding times of YAP and TEAD to control transcription. Sci Adv. 2025;11(30):eadw4974. Epub 20250725. doi: 10.1126/sciadv.adw4974. PubMed PMID: 40712036.

43. Jukam D, Xie B, Rister J, Terrell D, Charlton-Perkins M, Pistillo D, et al. Opposite feedbacks in the Hippo pathway for growth control and neural fate. Science. 2013;342(6155):1238016. Epub 2013/08/31. doi: 10.1126/science.1238016. PubMed PMID: 23989952; PubMed Central PMCID: PMC3796000.

44. Pojer JM, Manning SA, Kroeger B, Kondo S, Harvey KF. The Hippo pathway uses different machinery to control cell fate and organ size. iScience. 2021;24(8):102830. Epub 2021/08/07. doi: 10.1016/j.isci.2021.102830. PubMed PMID: 34355153; PubMed Central PMCID: PMCPMC8322298.

45. Nishioka N, Inoue K, Adachi K, Kiyonari H, Ota M, Ralston A, et al. The Hippo signaling pathway components Lats and Yap pattern Tead4 activity to distinguish mouse trophectoderm from inner cell mass. Dev Cell. 2009;16(3):398–410. Epub 2009/03/18. doi: S1534-5807(09)00077-X [pii] 10.1016/j.devcel.2009.02.003 [doi]. PubMed PMID: 19289085.

46. Home P, Saha B, Ray S, Dutta D, Gunewardena S, Yoo B, et al. Altered subcellular localization of transcription factor TEAD4 regulates first mammalian cell lineage commitment. Proc Natl Acad Sci U S A. 2012;109(19):7362–7. Epub 20120423. doi: 10.1073/pnas.1201595109. PubMed PMID: 22529382; PubMed Central PMCID: PMCPMC3358889.

47. Nicolay BN, Bayarmagnai B, Moon NS, Benevolenskaya EV, Frolov MV. Combined inactivation of pRB and hippo pathways induces dedifferentiation in the Drosophila retina. PLoS genetics. 2010;6(4):e1000918. Epub 2010/04/28. doi: 10.1371/journal.pgen.1000918. PubMed PMID: 20421993; PubMed Central PMCID: PMCPMC2858677.

48. Kowalczyk W, Romanelli L, Atkins M, Hillen H, Bravo Gonzalez-Blas C, Jacobs J, et al. Hippo signaling instructs ectopic but not normal organ growth. Science. 2022;378(6621):eabg3679. Epub 20221118. doi: 10.1126/science.abg3679. PubMed PMID: 36395225.

49. Fletcher GC, Diaz-de-la-Loza MD, Borreguero-Munoz N, Holder M, Aguilar-Aragon M, Thompson BJ. Mechanical strain regulates the Hippo pathway in Drosophila. Development. 2018. Epub 2018/02/15. doi: 10.1242/dev.159467. PubMed PMID: 29440303.

50. Friesen S, Hariharan IK. Coordinated growth of linked epithelia is mediated by the Hippo pathway. bioRxiv. 2023. Epub 20230317. doi: 10.1101/2023.02.26.530099. PubMed PMID: 36993542; PubMed Central PMCID: PMCPMC10054945.

51. Burke R, Basler K. Hedgehog signaling in Drosophila eye and limb development - conserved machinery, divergent roles? Curr Opin Neurobiol. 1997;7(1):55–61. doi: 10.1016/s0959-4388(97)80120-1. PubMed PMID: 9039793.

52. Huang J, Wu S, Barrera J, Matthews K, Pan D. The Hippo signaling pathway coordinately regulates cell proliferation and apoptosis by inactivating Yorkie, the Drosophila Homolog of YAP. Cell. 2005;122(3):421–34. PubMed PMID: 16096061.

53. Rodriguez A, Oliver H, Zou H, Chen P, Wang X, Abrams JM. Dark is a Drosophila homologue of Apaf-1/CED-4 and functions in an evolutionarily conserved death pathway. Nat Cell Biol. 1999;1(5):272–9. doi: 10.1038/12984. PubMed PMID: 10559939.

54. Mills K, Daish T, Harvey KF, Pfleger CM, Hariharan IK, Kumar S. The Drosophila melanogaster Apaf-1 homologue ARK is required for most, but not all, programmed cell death. J Cell Biol. 2006;172(6):809–15. PubMed PMID: 16533943.

55. Koontz LM, Liu-Chittenden Y, Yin F, Zheng Y, Yu J, Huang B, et al. The Hippo effector Yorkie controls normal tissue growth by antagonizing scalloped-mediated default repression. Dev Cell. 2013;25(4):388–401. Epub 2013/06/04. doi: 10.1016/j.devcel.2013.04.021. PubMed PMID: 23725764; PubMed Central PMCID: PMC3705890.

56. Vissers JHA, Dent LG, House CM, Kondo S, Harvey KF. Pits and CtBP Control Tissue Growth in Drosophila melanogaster with the Hippo Pathway Transcription Repressor Tgi. Genetics. 2020;215(1):117–28. Epub 2020/03/04. doi: 10.1534/genetics.120.303147. PubMed PMID: 32122936; PubMed Central PMCID: PMCPMC7198276.

57. Higashijima S, Kojima T, Michiue T, Ishimaru S, Emori Y, Saigo K. Dual Bar homeo box genes of Drosophila required in two photoreceptor cells, R1 and R6, and primary pigment cells for normal eye development. Genes Dev. 1992;6(1):50–60. doi: 10.1101/gad.6.1.50. PubMed PMID: 1346120.

58. Hayashi T, Kojima T, Saigo K. Specification of primary pigment cell and outer photoreceptor fates by BarH1 homeobox gene in the developing Drosophila eye. Dev Biol. 1998;200(2):131–45. PubMed PMID: 9705222.

59. Meserve JH, Duronio RJ. A population of G2-arrested cells are selected as sensory organ precursors for the interommatidial bristles of the Drosophila eye. Dev Biol. 2017;430(2):374–84. Epub 20170621. doi: 10.1016/j.ydbio.2017.06.023. PubMed PMID: 28645749; PubMed Central PMCID: PMCPMC5623626.

60. Fu W, Duan H, Frei E, Noll M. shaven and sparkling are mutations in separate enhancers of the Drosophila Pax2 homolog. Development. 1998;125(15):2943–50. doi: 10.1242/dev.125.15.2943. PubMed PMID: 9655816.

61. Blochlinger K, Jan LY, Jan YN. Postembryonic patterns of expression of cut, a locus regulating sensory organ identity in Drosophila. Development. 1993;117(2):441–50. doi: 10.1242/dev.117.2.441. PubMed PMID: 8330519.

62. Lee T, Luo L. Mosaic analysis with a repressible cell marker for studies of gene function in neuronal morphogenesis. Neuron. 1999;22(3):451–61. PubMed PMID: 10197526.

63. Perkins LA, Holderbaum L, Tao R, Hu Y, Sopko R, McCall K, et al. The Transgenic RNAi Project at Harvard Medical School: Resources and Validation. Genetics. 2015;201(3):843–52. Epub 20150828. doi: 10.1534/genetics.115.180208. PubMed PMID: 26320097; PubMed Central PMCID: PMCPMC4649654.

64. Dietzl G, Chen D, Schnorrer F, Su KC, Barinova Y, Fellner M, et al. A genome-wide transgenic RNAi library for conditional gene inactivation in Drosophila. Nature. 2007;448(7150):151–6. PubMed PMID: 17625558.

65. Mitchell KA, Vissers JHA, Pojer JM, Brooks E, Hilmi AJS, Papenfuss AT, et al. The JNK and Hippo pathways control epithelial integrity and prevent tumor initiation by regulating an overlapping transcriptome. Curr Biol. 2024;34(17):3966–82 e7. Epub 20240814. doi: 10.1016/j.cub.2024.07.060. PubMed PMID: 39146938.

66. Lai ZC, Rubin GM. Negative control of photoreceptor development in Drosophila by the product of the yan gene, an ETS domain protein. Cell. 1992;70(4):609–20. doi: 10.1016/0092-8674(92)90430-k. PubMed PMID: 1505027.

67. Halder G, Johnson RL. Hippo signaling: growth control and beyond. Development. 2011;138(1):9–22. Epub 2010/12/09. doi: 138/1/9 [pii] 10.1242/dev.045500. PubMed PMID: 21138973.

68. Xu T, Rubin GM. Analysis of genetic mosaics in developing and adult Drosophila tissues. Development. 1993;117(4):1223–37. Epub 1993/04/01. doi: 10.1242/dev.117.4.1223. PubMed PMID: 8404527.

69. Olson ER, Pancratov R, Chatterjee SS, Changkakoty B, Pervaiz Z, DasGupta R. Yan, an ETS-domain transcription factor, negatively modulates the Wingless pathway in the Drosophila eye. EMBO Rep. 2011;12(10):1047–54. Epub 20110930. doi: 10.1038/embor.2011.159. PubMed PMID: 21869817; PubMed Central PMCID: PMCPMC3185344.

70. Jukam D, Desplan C. Binary regulation of Hippo pathway by Merlin/NF2, Kibra, Lgl, and Melted specifies and maintains postmitotic neuronal fate. Dev Cell. 2011;21(5):874–87. Epub 2011/11/08. doi: S1534-5807(11)00421-7 [pii] 10.1016/j.devcel.2011.10.004. PubMed PMID: 22055343.

71. Rader AE, Bayarmagnai B, Frolov MV. Combined inactivation of RB and Hippo converts differentiating Drosophila photoreceptors into eye progenitor cells through derepression of homothorax. Dev Cell. 2023;58(21):2261–74 e6. Epub 20231016. doi: 10.1016/j.devcel.2023.09.003. PubMed PMID: 37848027; PubMed Central PMCID: PMCPMC10842633.

72. Charlton-Perkins M, Cook TA. Building a Fly Eye: Terminal Differentiation Events of the Retina, Corneal Lens, and Pigmented Epithelia. In: Cagan RL, Reh TABTCTiDB, editors. Invertebrate and Vertebrate Eye Development. 93: Academic Press; 2010. p. 129–73.

73. Lim HY, Tomlinson A. Organization of the peripheral fly eye: the roles of Snail family transcription factors in peripheral retinal apoptosis. Development. 2006;133(18):3529–37. Epub 20060816. doi: 10.1242/dev.02524. PubMed PMID: 16914498.

74. Wei GH, Badis G, Berger MF, Kivioja T, Palin K, Enge M, et al. Genome-wide analysis of ETS-family DNA-binding in vitro and in vivo. EMBO J. 2010;29(13):2147–60. Epub 20100601. doi: 10.1038/emboj.2010.106. PubMed PMID: 20517297; PubMed Central PMCID: PMCPMC2905244.

75. Wang H, Bollepogu Raja KK, Yeung K, Morrison CA, Terrizzano A, Khodadadi-Jamayran A, et al. Synergistic activation by Glass and Pointed promotes neuronal identity in the Drosophila eye disc. Nat Commun. 2024;15(1):7091. Epub 20240817. doi: 10.1038/s41467-024-51429-z. PubMed PMID: 39154080; PubMed Central PMCID: PMCPMC11330500.

76. Bernasek SM, Hur SSJ, Pelaez-Restrepo N, Boisclair Lachance JF, Bakker R, Navarro HT, et al. Ratiometric sensing of Pnt and Yan transcription factor levels confers ultrasensitivity to photoreceptor fate transitions in Drosophila. Development. 2023;150(8). Epub 20230424. doi: 10.1242/dev.201467. PubMed PMID: 36942737; PubMed Central PMCID: PMCPMC10163347.

77. Harvey KF, Tang TT. Targeting the Hippo pathway in cancer. Nature Reviews Drug Discovery. 2025. doi: 10.1038/s41573-025-01234-0.

78. Zanconato F, Forcato M, Battilana G, Azzolin L, Quaranta E, Bodega B, et al. Genome-wide association between YAP/TAZ/TEAD and AP-1 at enhancers drives oncogenic growth. Nature Cell Biology. 2015;17(9):1218–27. doi: 10.1038/ncb3216.

79. Pascual J, Jacobs J, Sansores-Garcia L, Natarajan M, Zeitlinger J, Aerts S, et al. Hippo Reprograms the Transcriptional Response to Ras Signaling. Dev Cell. 2017;42(6):667–80 e4. Epub 2017/09/28. doi: 10.1016/j.devcel.2017.08.013. PubMed PMID: 28950103.

80. Kulkarni A, Mohan V, Tang TT, Post L, Chan YC, Manning M, et al. Identification of resistance mechanisms to small-molecule inhibition of TEAD-regulated transcription. EMBO Rep. 2024;25(9):3944–69. Epub 20240805. doi: 10.1038/s44319-024-00217-3. PubMed PMID: 39103676; PubMed Central PMCID: PMCPMC11387499.

81. Hagenbeek TJ, Zbieg JR, Hafner M, Mroue R, Lacap JA, Sodir NM, et al. An allosteric pan-TEAD inhibitor blocks oncogenic YAP/TAZ signaling and overcomes KRAS G12C inhibitor resistance. Nature Cancer. 2023;4(6):812–28. doi: 10.1038/s43018-023-00577-0.

82. Paul S, Hagenbeek TJ, Tremblay J, Kameswaran V, Ong C, Liu C, et al. Cooperation between the Hippo and MAPK pathway activation drives acquired resistance to TEAD inhibition. Nat Commun. 2025;16(1):1743. Epub 20250218. doi: 10.1038/s41467-025-56634-y. PubMed PMID: 39966375; PubMed Central PMCID: PMCPMC11836325.

83. Edwards AC, Stalnecker CA, Jean Morales A, Taylor KE, Klomp JE, Klomp JA, et al. TEAD Inhibition Overcomes YAP1/TAZ-Driven Primary and Acquired Resistance to KRASG12C Inhibitors. Cancer Res. 2023;83(24):4112–29. doi: 10.1158/0008-5472.can-23-2994. PubMed PMID: 37934103; PubMed Central PMCID: PMCPMC10821578.

84. Mukhopadhyay S, Huang HY, Lin Z, Ranieri M, Li S, Sahu S, et al. Genome-Wide CRISPR Screens Identify Multiple Synthetic Lethal Targets That Enhance KRASG12C Inhibitor Efficacy. Cancer Res. 2023;83(24):4095–111. doi: 10.1158/0008-5472.can-23-2729. PubMed PMID: 37729426; PubMed Central PMCID: PMCPMC10841254.

85. Campbell SD, Duttaroy A, Katzen AL, Chovnick A. Cloning and characterization of the scalloped region of Drosophila melanogaster. Genetics. 1991;127(2):367–80. doi: 10.1093/genetics/127.2.367. PubMed PMID: 1706292; PubMed Central PMCID: PMCPMC1204364.

86. Genevet A, Polesello C, Blight K, Robertson F, Collinson LM, Pichaud F, et al. The Hippo pathway regulates apical-domain size independently of its growth-control function. J Cell Sci. 2009;122(Pt 14):2360–70. Epub 2009/06/18. doi: jcs.041806 [pii] 10.1242/jcs.041806. PubMed PMID: 19531586.

87. Vincent JP, Girdham CH, O’Farrell PH. A cell-autonomous, ubiquitous marker for the analysis of Drosophila genetic mosaics. Dev Biol. 1994;164(1):328–31. doi: 10.1006/dbio.1994.1203. PubMed PMID: 8026635.

88. Golic KG, Lindquist S. The FLP recombinase of yeast catalyzes site-specific recombination in the Drosophila genome. Cell. 1989;59(3):499–509. doi: 10.1016/0092-8674(89)90033-0. PubMed PMID: 2509077.

89. Golic KG. Site-specific recombination between homologous chromosomes in Drosophila. Science. 1991;252(5008):958–61. doi: 10.1126/science.2035025. PubMed PMID: 2035025.

90. Lee T, Luo L. Mosaic analysis with a repressible cell marker (MARCM) for Drosophila neural development. Trends Neurosci. 2001;24(5):251–4. PubMed PMID: 11311363.

91. Manning SA, Dent LG, Kondo S, Zhao ZW, Plachta N, Harvey KF. Dynamic Fluctuations in Subcellular Localization of the Hippo Pathway Effector Yorkie In Vivo. Curr Biol. 2018;28(10):1651–60 e4. Epub 2018/05/15. doi: 10.1016/j.cub.2018.04.018. PubMed PMID: 29754899.

92. Kroeger B, Manning SA, Fonseka Y, Oorschot V, Crawford SA, Ramm G, et al. Basal spot junctions of Drosophila epithelial tissues respond to morphogenetic forces and regulate Hippo signaling. Dev Cell. 2024;59(2):262–79 e6. Epub 20231221. doi: 10.1016/j.devcel.2023.11.024. PubMed PMID: 38134928.

93. Bao S, Cagan R. Preferential adhesion mediated by Hibris and Roughest regulates morphogenesis and patterning in the Drosophila eye. Dev Cell. 2005;8(6):925–35. doi: 10.1016/j.devcel.2005.03.011. PubMed PMID: 15935781.

94. Poon CL, Lin JI, Zhang X, Harvey KF. The Sterile 20-like Kinase Tao-1 Controls Tissue Growth by Regulating the Salvador-Warts-Hippo Pathway. Dev Cell. 2011;21(5):896–906. Epub 2011/11/15. doi: S1534-5807(11)00410-2 [pii] 10.1016/j.devcel.2011.09.012. PubMed PMID: 22075148.

95. Poon CLC, Liu W, Song Y, Gomez M, Kulaberoglu Y, Zhang X, et al. A Hippo-like Signaling Pathway Controls Tracheal Morphogenesis in Drosophila melanogaster. Dev Cell. 2018;47(5):564–75 e5. doi: 10.1016/j.devcel.2018.09.024. PubMed PMID: 30458981; PubMed Central PMCID: PMCPMC6281297.

96. Schindelin J, Arganda-Carreras I, Frise E, Kaynig V, Longair M, Pietzsch T, et al. Fiji: an open-source platform for biological-image analysis. Nat Methods. 2012;9(7):676–82. doi: 10.1038/nmeth.2019. PubMed PMID: 22743772; PubMed Central PMCID: PMCPMC3855844.

97. Charlton-Perkins M, Whitaker SL, Fei Y, Xie B, Li-Kroeger D, Gebelein B, et al. Prospero and Pax2 combinatorially control neural cell fate decisions by modulating Ras- and Notch-dependent signaling. Neural development. 2011;6(1):20. Epub 20110503. doi: 10.1186/1749-8104-6-20. PubMed PMID: 21539742; PubMed Central PMCID: PMCPMC3123624.

98. Gresser AL, Gutzwiller LM, Gauck MK, Hartenstein V, Cook TA, Gebelein B. Rhomboid Enhancer Activity Defines a Subset of Drosophila Neural Precursors Required for Proper Feeding, Growth and Viability. PLoS One. 2015;10(8):e0134915. Epub 20150807. doi: 10.1371/journal.pone.0134915. PubMed PMID: 26252385; PubMed Central PMCID: PMCPMC4529294.

